# Phase diversity improves retinal image quality in adaptive optics scanning light ophthalmoscopy

**DOI:** 10.64898/2025.12.16.694208

**Authors:** Yongyi Cai, Eryk Druszkiewicz, Sara S. Patterson, Keith Parkins, Juliette E. Mcgregor, William H. Merigan, James R. Fienup, David R. Williams

## Abstract

The quality of retinal images is compromised by aberrations that remain uncorrected even in confocal adaptive optics imaging. This study demonstrates phase diversity (PD), a computational imaging technique, to address residual aberrations and enhance image quality in adaptive optics scanning laser ophthalmoscopy (AOSLO). By using images of the same object obtained with and without deliberately added aberrations, PD computes and compensates for the effects of existing residual aberrations beyond those corrected by a closed-loop AO system. Experimental validation demonstrates that PD improves visualization of retinal microstructures, including cone mosaics and dendrites of fluorescently labeled retinal ganglion cells (RGCs).

## 1. Introduction

Adaptive optics scanning laser ophthalmoscopy (AOSLO) provides non-invasive imaging of the retina at cellular resolution by compensating for the optical aberrations, which has significantly advanced the understanding of retinal structure and function [1–3]. However, the performance of AOSLO remains constrained by residual aberrations that persist even after hardware-based AO correction. These residual errors may arise from non-common path aberrations (NCPAs)— introduced in the detection path but not sampled by the wavefront sensor and thus left uncorrected by the deformable mirror [4,5]—or from uncorrected dynamic aberrations arising from tear film fluctuations, intraocular scattering, and physiological instabilities such as respiration and heartbeat, which exceed the finite temporal correction bandwidth of the adaptive optics system [6,7]. In addition, AO performance is limited by the finite spatial bandwidth set by the Shack–Hartmann sampling and deformable mirror actuator density. Although modern ophthalmic AO systems can readily correct Zernike modes beyond the 7th radial order, still higher-order aberrations lie outside this spatial correction range and therefore remain uncorrected [8,9]. Such residual aberrations reduce image contrast and resolution, limiting the visibility of fine subcellular structures in the retina. To mitigate this problem, we investigated phase diversity as a complementary method to recover and correct residual static aberrations beyond those compensated by closed-loop AO.

Originally introduced for astronomical imaging to compensate for atmospheric turbulence, phase diversity (PD) jointly estimates the object and aberrated wavefront from two (or more) images acquired with known phase differences, typically one in-focus and one slightly out-of-focus [10–12]. By fitting images computed from a forward model of the imaging system to measured data, PD simultaneously reconstructs both the object and the aberrations, offering improved robustness to noise. Variants of PD have been adapted for widefield microscopy to effectively correct static, sample-induced aberrations in extended biological specimens [13–16]. However, it has not been applied to the eye with AO confocal scanning. In the confocal scanning imaging modality, illumination and detection are spatially localized through a scanning beam and a confocal pinhole [17,18]. This confocal configuration fundamentally alters the forward model typically assumed in widefield PD approaches. Additionally, sequential scanning in AOSLO introduces temporal variability [19–21] absent in widefield microscopy, complicating the direct transfer of existing PD methods. Thus, applying PD to AOSLO imaging requires a confocal imaging forward model and the associated inversion algorithms to accommodate confocality and scanning-related artifacts. These considerations raise some questions:

1. Can PD techniques be effectively adapted to accommodate the confocal and scanning geometry of AOSLO?
2. How accurately can PD algorithms recover residual wavefront errors from realistic, noise-affected, and motion-impacted AOSLO image pairs?
3. What quantitative enhancements in resolution, contrast, and point-spread function (PSF) fidelity can PD bring to AOSLO images beyond conventional AO hardware corrections? In this paper, we validated a Zernike-polynomial-based PD algorithm specifically tailored for confocal AOSLO imaging. We assessed its performance in wavefront-sensing accuracy and image quality using static model eyes and *in vivo* retinal images. To the best of our knowledge, this work represents the first successful integration of PD with confocal AOSLO post-processing for robust correction of residual aberrations, thereby significantly enhancing image quality beyond the capabilities of standard AO techniques alone.

## 2. Methods

### 2.1 Animal care

Two macaque monkeys (Macaca fascicularis), both males, were used in this study. The animals were housed in pairs in an AAALAC-accredited facility under the care of the University of Rochester’s Department of Comparative Medicine. Their daily care was provided by a team including multiple full-time veterinarians, veterinary technicians, an animal behaviorist, and animal care staff who monitored health and comfort at least twice daily. The macaques had free access to water and a complete diet of primate biscuits, supplemented with a variety of treats such as nuts, dried and fresh fruits, vegetables, vitamin chews, and high-value rewards including marshmallows and fruit gummies. Enrichment was provided daily and included mirrors, puzzle feeders, durable chew toys, as well as television or music sessions. Novel enrichment items, such as treat-filled forage boxes, grapevines, water play, and natural materials like grass or snow, were introduced multiple times weekly. The animals also had rotating access to a large play space equipped with swings and perches. All experiments were performed in strict accordance with the Association for Research in Vision and Ophthalmology (ARVO) Statement for the Use of Animals in Ophthalmic and Vision Research and the recommendations in the Guide for the Care and Use of Laboratory Animals of the National Institutes of Health. The experimental protocol was approved by the University Committee on Animal Resources (UCAR) of the University of Rochester (PHS assurance number: D16-00188 (A3292-01)).

### 2.2 Intravitreal injections

Intravitreal delivery of adeno-associated viral (AAV2) vectors was performed under sterile conditions as previously described [22]. The ocular surface was disinfected with diluted povidone-iodine prior to injection. A tuberculin syringe fitted with a 30-gauge needle was used to deliver the viral suspension into the mid-vitreous through the pars plana, approximately 2– 4 mm posterior to the limbus. Injections consisted of 75-100 *μL* of vector solution, including *AAV2–CaMKIIa-mCherry* (Addgene) in OD of M1 and *AAV2-hSyn1-ChRmine-mScarlet* (Shannon and Sanford Boye Laboratory, University of Florida). To minimize immune responses, animals received daily subcutaneous cyclosporine A, with blood trough levels maintained between 150-200 *ng*/*mL*, and intravitreal triamcinolone was administered following injection when indicated. Expression and ocular health were monitored with scanning light ophthalmoscopy (Spectralis HRA, Heidelberg Engineering) using the 488 nm autofluorescence mode on a regular basis.

### 2.3 Imaging preparation

All animal procedures were carried out under the supervision of licensed veterinary technicians in compliance with institutional and federal guidelines. Prior to each imaging session, macaques were fasted overnight and subsequently anesthetized with isoflurane on the morning of the procedure. Animals were positioned prone on a custom stereotaxic cart, stabilized with ear bars and a chin rest, cushioned for comfort, and maintained on a warming system to preserve body temperature. To ensure immobilization during imaging, vecuronium bromide was administered, and vital signs—including heart rate, blood pressure, respiration, oxygen saturation, and electrocardiogram—were continuously monitored and documented at regular intervals. Pupils were dilated using 2.5% phenylephrine and 1% tropicamide, and a rigid gas-permeable contact lens was applied to protect corneal hydration and improve optical quality. The stereotaxic cart allowed adjustment of body orientation to optimize eye positioning for imaging. Following each session, animals were closely monitored by veterinary staff until full recovery from anesthesia was confirmed.

### 2.4 Adaptive optics scanning laser ophthalmoscopy

Experiments were performed on a custom-built AOSLO optimized for high-resolution confocal retinal imaging that was described previously [23,24]. A Shack-Hartmann wavefront sensor, paired with an 847 nm diode laser beacon (QPhotonics), measured the ocular aberrations in real time. These data drove a 97-actuator deformable mirror (ALPAO) operating in closed loop at ~8 Hz, providing continuous aberration correction across a 7.2 mm pupil. The cone mosaic was visualized using a 796 nm superluminescent diode (Superlum), with reflected light collected through a 1.1 Airy Unit (AU, 20 *μm*) confocal pinhole. This channel served both to navigate to consistent retinal locations across sessions and to enable high-resolution strip-based motion correction of the fluorescent imaging videos. Retinal ganglion cells (RGCs) expressing mCherry and mScarlet were imaged using a 561 nm laser (Toptica), with emitted fluorescence collected through a pinhole at 2.5 AU. Emission was filtered at 607/40 nm for mCherry and 590/20 nm for mScarlet separately to collect at the highest emission range. All data were collected over a small field of view of either 1.09° × 1.09°or 0.81° × 0.81°and covered 496 × 496 pixels. This ensured that the pixel size (~0.1-0.12 arcmin) met the Nyquist sampling criterion (~0.175 arcmin) for the system’s theoretical diffraction-limited resolution (~ 0.35 Rayleigh angular resolution arcmin @590nm with 7.2mm pupil). Two consecutive PD images of the same retinal location were obtained by summing 750 frames for each, one with and one without a deliberately introduced defocus. These aberrations were introduced using a custom-built AO software that introduced the corresponding Zernike mode to the deformable mirror as a fixed bias outside the AO closed loop. All acquired frames were registered to correct for eye motion and averaged to improve the signal-to-noise ratio (SNR).

### 2.5 Phase diversity algorithm

Let *W*(*u*) denote the wavefront aberration over the normalized pupil, parameterized by Zernike coefficients. The coherent amplitude impulse-response (IPR) function is given by [25]

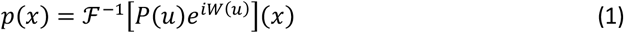

where *P*(*u*) is the pupil aperture function, and ℱ^−1^ denotes the inverse Fourier transform, and the intensity point-spread function (PS) is given by |*p*(*x*)|^2^ for a flood-illuminated system. For a confocal imaging system with a finite pinhole, the intensity point-spread function (PSF) is written as [26]

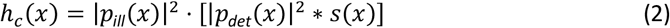

where

- *p*_*ill*_(*x*) and *p*_*det*_(*x*) are the illumination and detection IPRs, respectively, obtained from the inverse Fourier transform of the aberrated pupil,
- *s*(*x*) is the pinhole aperture function given by

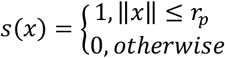

with *r*_*p*_ determined by the pinhole radius in AU, and
- ∗ denotes the 2D convolution.

Given two or more images acquired at different known phase diversities *ΔW*_*k*_, the forward model becomes

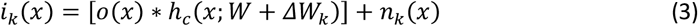

where

- *o*(*x*) is the object intensity-reflectivity distribution in the retina,
- *h*_*c*_(*x*; *W* + *ΔW*_*k*_) is the confocal PSF under the combined unknown aberration *W* and known aberrations *ΔW*_*k*_, and
- *n*_*k*_(*x*) represents photon noise and electronic noise. The wavefront *W* was parameterized by a Zernike expansion

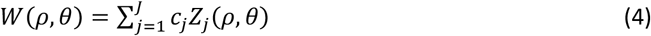

where *c*_j_ are the coefficients to be estimated. The inverse problem was formulated as minimizing the residual under the assumption of additive Gaussian read noise

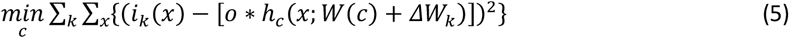

which corresponds to the maximum-likelihood estimator for Gaussian noise. Using Parseval’s theorem, the problem is equivalently expressed in the Fourier domain

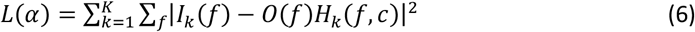

where *f* are Fourier domain spatial frequency indices, and *I, O* and *H* are the Fourier transforms of *i, o* and *h*, respectively. Notice that *H* is the optical transfer function (OTF) of the system, which depends explicitly on *c*, the unknown phase parameters. Note that *L* is a scalar objective function obtained by summing over all frequencies. For the case of two diversity images, given a fixed aberration function, a closed form estimate of the object spectrum can be obtained using a maximum likelihood estimator:

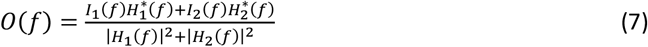

where the asterisk denotes complex conjugation. Substituting this expression for *O*(*f*) into Eq. (6) yields a reduced Gaussian log-likelihood cost to [11]

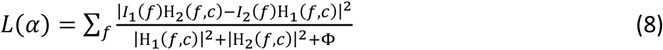

where we added a constant regularization parameter Φ = 10^−18^ chosen by the simulation demonstrated in **Fig. 1**. The phase was iteratively updated by minimizing *L*(*a*) in Eq. (8) until convergence.

**Fig. 1.**
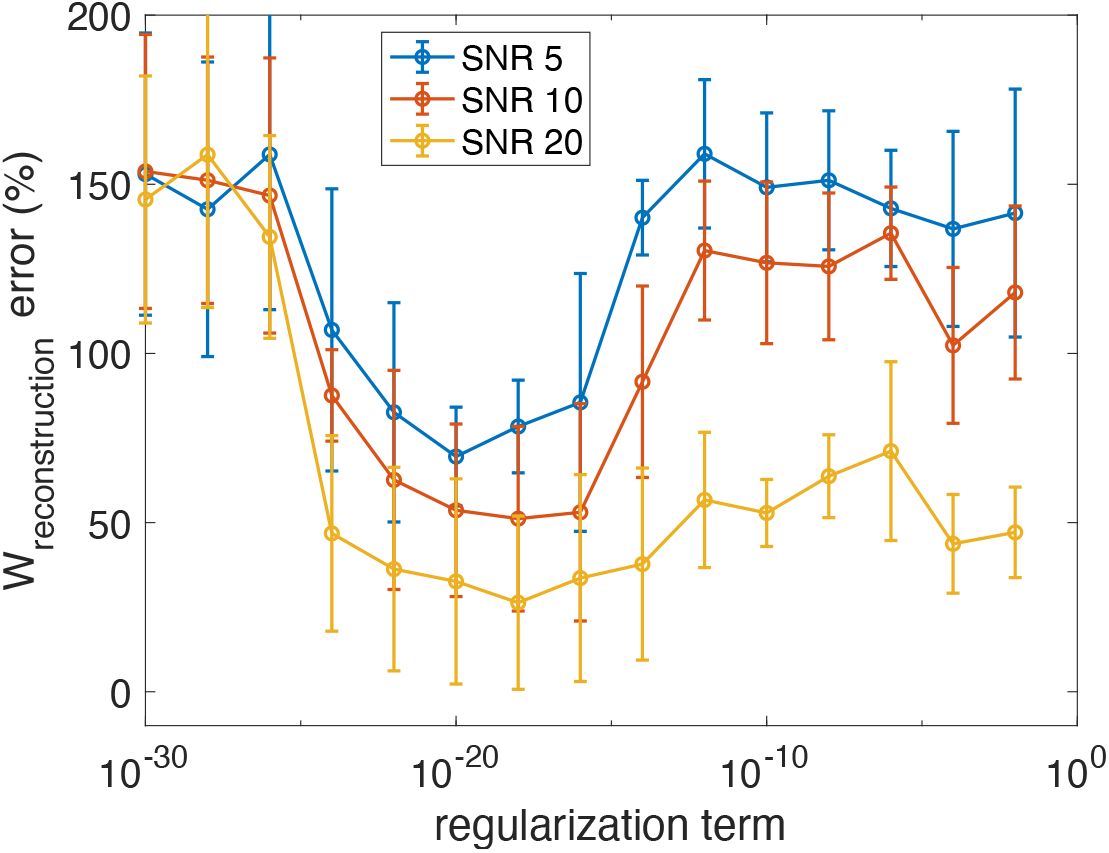
Effect of the regularization parameter on phase diversity (PD) performance. Wavefront reconstruction error is plotted as a function of the regularization term for different signal-to-noise ratios (SNR = 5, 10, 20). At very small regularization values, noise amplification degraded performance, while overly large values over-smoothed the solution. An intermediate range (10^−22^ - 10^−16^) provided stable and accurate reconstruction, with lower errors achieved at higher SNR. Error bars indicate standard deviations across repeated trials.

## 3. Results

### 3.1. Simulation of phase diversity performance on synthetic AOSLO data

To validate the phase diversity (PD) algorithm’s theoretical efficacy, we first tested it on synthetic AOSLO data with known aberrations. A computational model of the primate’s eye was generated, incorporating the morphology of a RGC taken from histology [27]. The pupil size used was 7.2 mm, and the focal length was 17 mm. The original image is composed of 632×632 pixels with a 0.2 *μm* pixel size at an image wavelength of 590 *nm*. The model incorporates background light, scaled appropriately relative to measurements of RGC fluorescence. An estimate of the background light was obtained from a through-focus measurement in a real eye that quantified the background fluorescence mainly from the nerve fiber layer (NFL) and the retinal pigment epithelium (RPE), which reduces the contrast of the signal originating from with labelled RGC. The measured photon noise and detector noise have also been incorporated [28]. Aberrations were introduced using Zernike polynomials (modes 3–15, OSA/ANSI index), having RMS = 0.149 waves @590 *nm w*avelength) to simulate ocular residual wavefront errors. The PD algorithm estimated the phase aberrations from pairs consisting of an in-focus image and an image with 0.8 PV-waves of astigmatism, iteratively minimizing the cost function.

**Fig. 2**. illustrates the performance of PD on a simulated retinal image. As shown in **Fig. 2(A)**, the presence of ocular aberrations substantially blurs fine dendritic structure, and the addition of 0.8 waves PV astigmatism further reduces contrast and structure sharpness. PD-based reconstruction restores much of this lost spatial detail: the reconstructed image is sharper and exhibits higher local contrast than either of the aberrated inputs. When combined with deconvolution, the reconstruction achieves spatial detail comparable to the unaberrated reference image acquired through the same finite pupil.

**Fig. 2.**
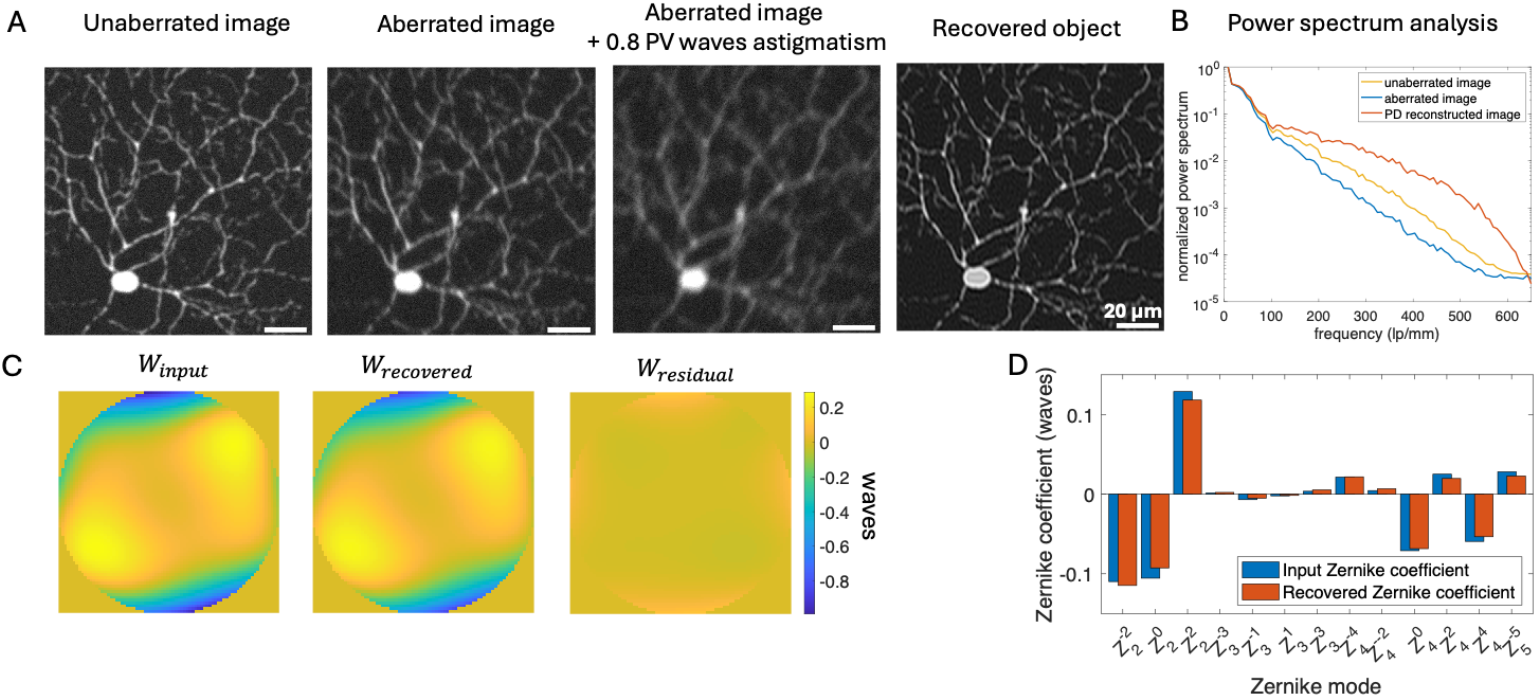
Simulation results validating phase diversity wavefront reconstruction. (A) Representative simulation showing the unaberrated reference image (no aberrations, but imaged through the same aperture), an aberrated image, the same aberrated image with additional 0.8 PV waves astigmatism (used as PD input), and the recovered image after PD correction. Scale bars = 20 *μm*. (B) Power spectrum analysis comparing the unaberrated, aberrated, and PD-recovered images. The PD-corrected reconstruction recovers high spatial-frequency content approaching the diffraction limit. (C) Wavefront maps showing the input (ground-truth) aberration, the recovered wavefront estimated by PD, and the residual wavefront. (D) Comparison of Zernike mode coefficients between the input and recovered wavefronts. Blue bars indicate ground truth; red bars indicate recovered values. The strong agreement across modes highlights the accuracy of PD in recovering high-order aberrations.

This improvement is quantitatively supported in **Fig. 2(B)**, where the power spectrum of the PD-corrected image recovers high-spatial-frequency content that had been suppressed by the aberration. The recovered aberration closely matched the input wavefront, with only a small residual error **(Fig. 2(C)**. The accuracy of Zernike recovery was further confirmed by the strong agreement between the input and recovered coefficients **(Fig. 2(D))**. Overall, the RMS wavefront error was reduced by 88% (from 0.222 waves to 0.026 waves RMS), demonstrating the ability of PD to accurately estimate and correct higher-order aberrations.

We further investigated the role of confocal pinhole size in shaping the performance of phase diversity. **Fig. 3**. summarizes the results for both astigmatism **(Fig. 3(A))** and defocus **(Fig. 3(B))** over a range of added aberration amplitudes. In both cases, PD accuracy was strongly influenced by the confocal geometry. With small pinholes (0.5 AU), reconstruction errors were high because the pinhole rejected much of the aberrated light, reducing sensitivity to wavefront errors. Intermediate sizes (1.5–2.5 AU) improved accuracy, while the largest pinhole (7.5 AU) consistently gave the best performance. This highlights the trade-off when applying phase diversity to confocal images whereas small pinholes sharpen confocal images by filtering out aberrations which PD requires to be preserved, making larger pinholes more effective for wavefront recovery. While recovering the full wavefront with larger confocal pinholes improves image contrast and lateral resolution, smaller pinholes intrinsically provide superior lateral resolution [29]. Thus, the optimal pinhole diameter depends on whether image quality is primarily limited by residual aberrations or by the confocal aperture itself. The simulations presented here primarily isolate the effect of pinhole size on wavefront reconstruction under high-SNR, 2D conditions. However, a larger pinhole also increases photon throughput, reducing the typical trade-off associated with confocal optical sectioning. Instead of discarding out-of-focus light, PD can leverage this additional signal to improve the wavefront estimate. In this regime, the higher photon budget can tip the balance in favor of PD—especially in weak-fluorescence applications where photons rejected by a small pinhole cannot be recovered.

**Fig. 3.**
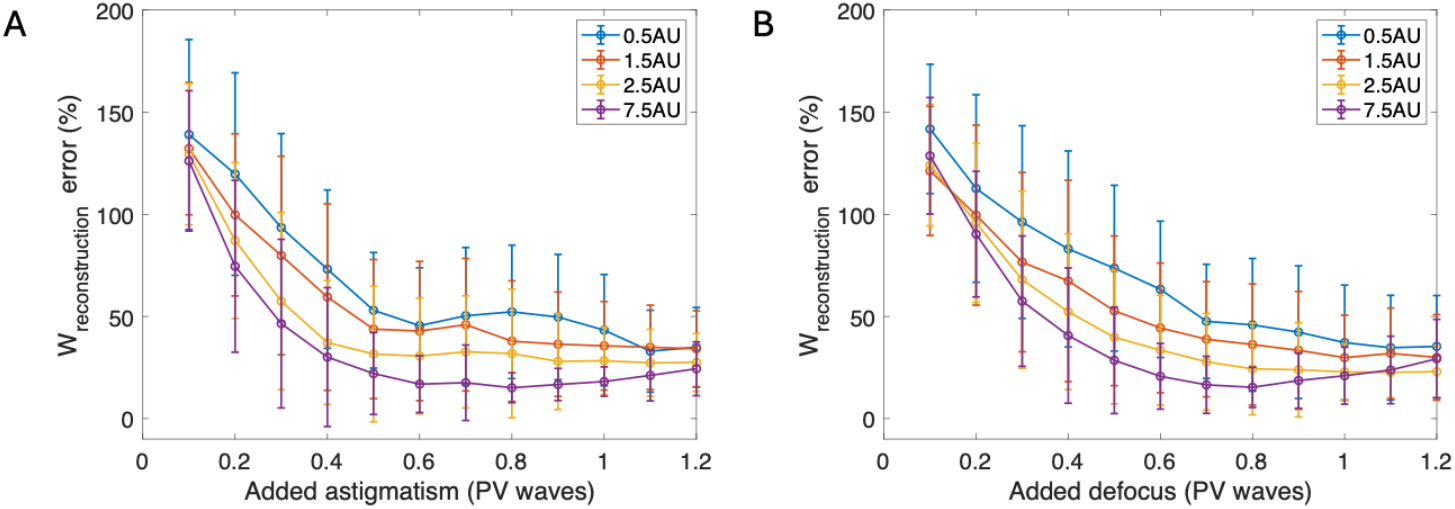
Effect of confocal pinhole size on phase diversity (PD) performance in AOSLO. (A) Wavefront RMS error as a function of added astigmatism for different confocal pinhole diameters (0.5–7.5 Airy units, AU). (B) Corresponding curves for added defocus. Means and standard deviations from 50 simulations are shown. In both cases, PD reconstruction accuracy depends strongly on the confocal pinhole size. Larger pinhole sizes and moderate diversity amplitudes improved reconstruction accuracy.

#### 3.2. Validation using a model eye and known phase

To quantify PD’s accuracy under controlled experimental conditions, we introduced calibrated aberrations (RMS = 0.31 waves) into a model eye system using a deformable mirror (DM). The model eye system consisted of a 7mm diameter pupil aperture and a 19mm focal-length focusing lens, mimicking the primate’s eye optics, and a piece of paper as the model of the retina, with a 7.5 AU pinhole. The AOSLO operated in an open-loop configuration during the test, allowing the Shack–Hartmann wavefront sensor to directly record the wavefront imposed by the DM. These measurements were later converted into wavefront maps and decomposed into Zernike coefficients to serve as the known ground truth. Reflectance images were acquired from the model eye both with and without the imposed aberrations. Subsequently, PD was applied to the aberrated image set to recover the wavefront distortions.

The results, summarized in **Fig. 4**., demonstrate the efficacy of PD in accurately retrieving and compensating aberrations. **Fig. 4(A)** shows images from the model eye experiment, including the unaberrated model eye structure, the aberrated image introduced by the deformable mirror (DM), an additional degradation with 0.97 PV waves of astigmatism, and the PD-reconstructed image. Following PD correction, the reconstructed image closely resembled the original unaberrated case, with fine features restored. This recovery is reflected in the frequency domain (**Fig. 4(B)**), where the aberrated image shows a substantial loss of mid- and high-spatial-frequency content that is restored following PD reconstruction.

**Fig. 4.**
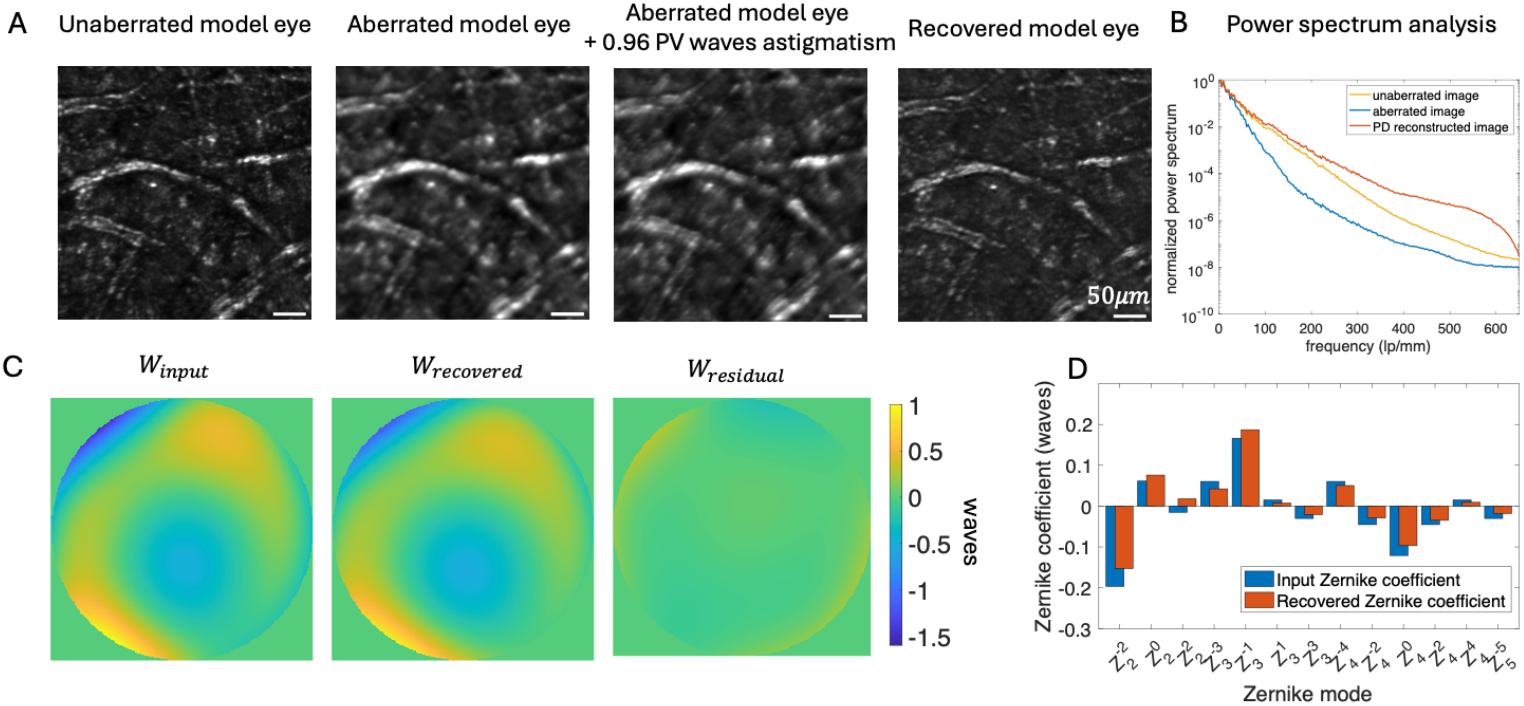
Model eye results validating phase diversity wavefront reconstruction. (A) Reflectance images acquired by the AOSLO: the unaberrated model eye, the same case with introduced aberrations (0.31 waves RMS), the aberrated case plus an additional 0.96 waves of astigmatism for PD acquisition, and the final PD-corrected reconstruction. Scale bars: 50*μm*. (B) Power spectrum comparison illustrating restoration of mid–high spatial frequencies after PD. (C) Wavefront maps showing the input wavefront applied by the deformable mirror, the PD-recovered wavefront, and the residual wavefront error (0.101 waves RMS). (D) Comparison of recovered (red) and ground-truth input (blue) Zernike coefficients, indicating accurate retrieval of the introduced aberrations.

Quantitatively, PD reduced the RMS wavefront error by 67% (from 0.31 waves to 0.1 waves), and most of the recovered Zernike coefficients closely matched the ground-truth values imposed by the DM (**Fig. 4(C) and (D)**), indicating accurate retrieval of the wavefront profile. These results validated the ability of PD to effectively estimate and correct aberrations in an AOSLO system, preserving diffraction-limited resolution even in the presence of imposed wavefront errors.

To evaluate the impact of confocal pinhole size on wavefront reconstruction, we imaged the same model-eye region while introducing known aberrations across two pinhole diameters in 4 trials (**Fig. 5(A)**). With a 7.5 AU pinhole, the PD-estimated astigmatism amplitude closely matched the ground-truth input of 1.07 PV waves (1.10 ± 0.22 PV, mean ± SD, **Fig. 5(B)**). In contrast, using a smaller 2.5 AU pinhole reduced the collected signal and degraded the reliability of the wavefront retrieval, leading to systematic underestimation of the true aberration (0.77 ± 0.12 PV, mean ± SD, **Fig. 5(B)**). This underestimation trend was consistent across repeated trials and agreed with our simulation results, although the experimental effect was even more pronounced. In simulation, we primarily attributed this bias to the reduced transmission of high spatial frequencies by smaller pinholes, which attenuates the blur induced by aberrations PD uses as a cue for phase retrieval. Experimentally, the effect is amplified by two other factors: (1) reduced photon throughput (lower SNR) and (2) increased optical sectioning, which changes the axial weighting of structures and thus the observed blur signature. Together, these factors diminish PD’s sensitivity to the true aberration when the pinhole is too small.

**Fig. 5.**
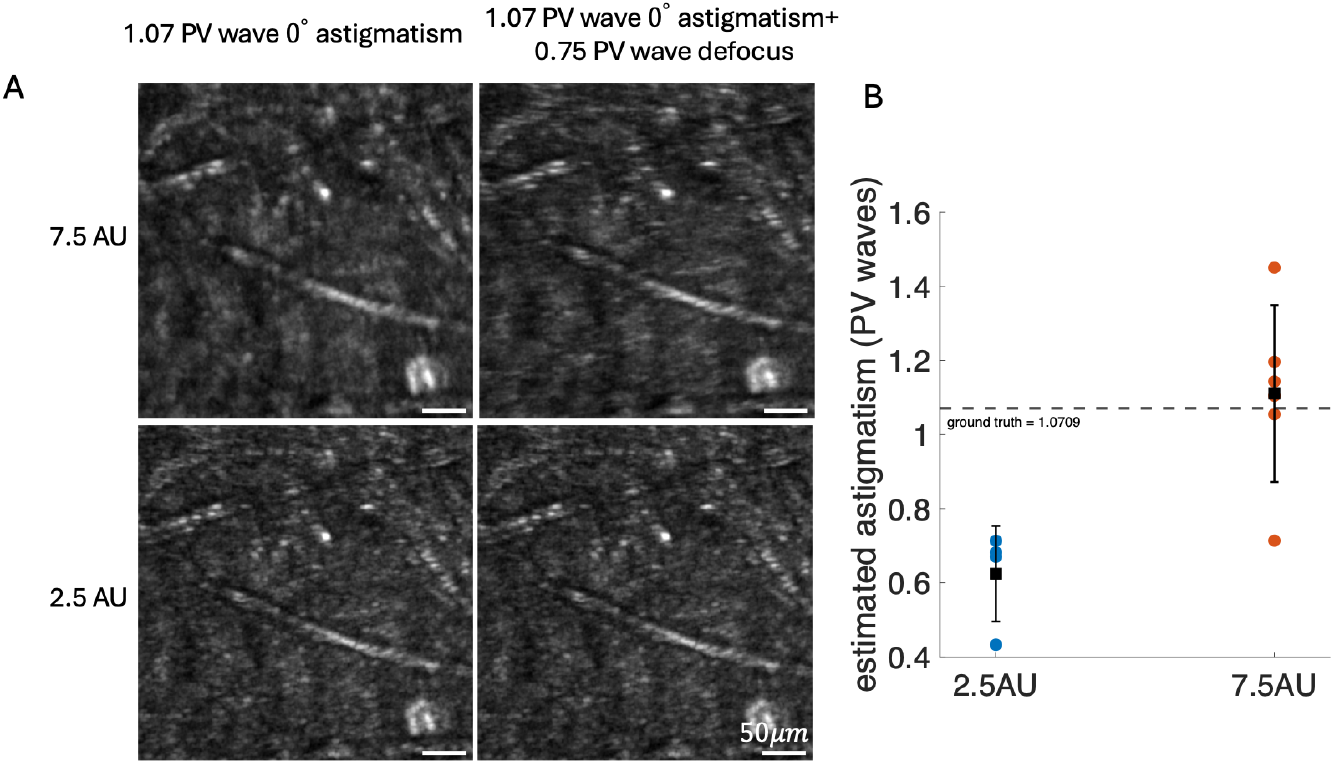
Wavefront reconstruction under different confocal pinhole sizes. (A) Example of AOSLO images of the same model eye region with added aberrations. Left: 1.07 PV waves of 0° astigmatism. Right: 1.07 PV waves of 0° astigmatism combined with 0.75 PV waves of defocus for 7.5 AU (top) and 2.5 AU (bottom) pinholes. Scale bar: 50 *μm*. (B) Quantification of the recovered astigmatism amplitudes under different pinhole sizes. Each dot represents an independent trial; black squares denote mean values with error bars showing ±1 standard deviation. The dashed line marks the ground-truth input astigmatism (1.07 PV waves). With a large pinhole (7.5 AU), the recovered astigmatism closely matched the ground truth. In contrast, smaller pinholes (2.5 AU) led to systematic underestimation of the aberration amplitude.

### 3.3. PD for *in vivo* monkey imaging data

To evaluate the performance of phase diversity for in vivo imaging of distinct retinal structures, we applied the PD algorithm to both reflectance and fluorescence AOSLO imaging collected from macaques.

**Fig. 6**. shows reflectance AOSLO images of the photoreceptor mosaic obtained from two macaques, each with two retinal regions. For each region, the defocused image, in-focus image, and PD-reconstructed image are displayed side by side. In both macaques, PD reconstruction recovers sharper cone boundaries and improves local contrast relative to the in-focus images, revealing finer details of the cone mosaic. The corresponding radial power spectra (right column) demonstrate an enhancement of high spatial frequencies following PD reconstruction, particularly at the spatial frequencies that correspond to the average cone spacing, where there is a local maximum in the power spectrum.

**Fig. 6.**
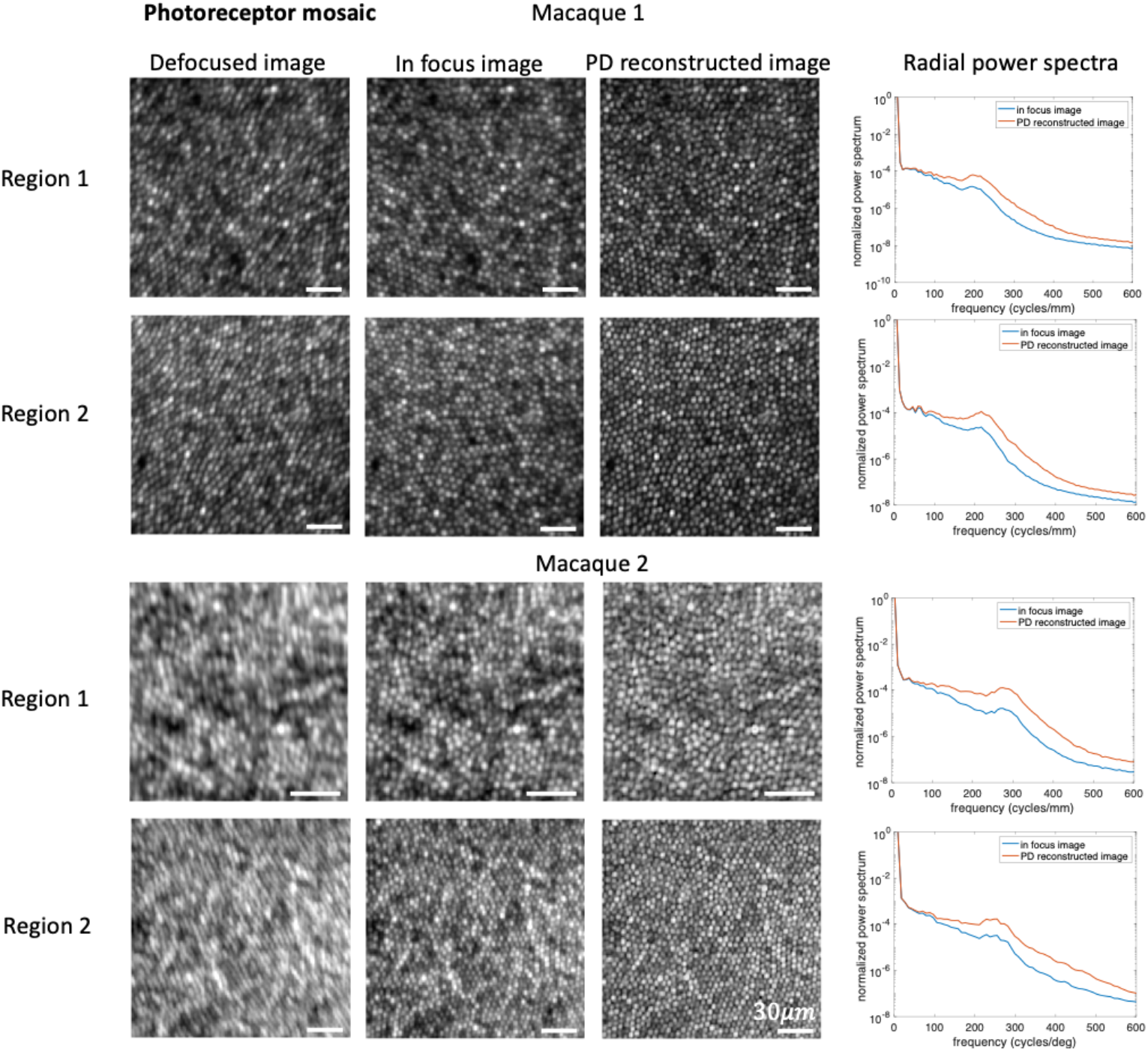
Reflectance AOSLO images of photoreceptors mosaic from two macaques, each with two retinal regions. For each region, the defocused image, in-focus image, and PD reconstructed image are shown side-by-side and in logarithmic scale. In both animals, PD reconstruction recovers sharper cone spacing and improves local contrast relative to the in-focus image. Corresponding radial power spectra (right column) demonstrate a substantial enhancement of high spatial frequencies following PD correction. Scale bars: 30*μm*.

Similarly, when applied to fluorescence images of RGCs, PD reconstruction enhanced the visualization of fine dendritic processes that appeared blurred or partially unresolved in the in-focus images (**Fig. 7.)**. Quantitative analysis of cross-sectional intensity profiles from 30 dendrites across multiple locations demonstrated an average 16% ± 8% decrease in full-width half-maximum (FWHM) (**Fig. 8**.), indicating significant sharpening of structural details. Moreover, the combination of lower background noise and enhanced signal intensity increased the signal-to-noise ratio (SNR) by approximately a factor of 1.5. Importantly, the PD results were generated using the same number of frames as the corresponding in-focus images, ensuring that the observed SNR improvement did not arise from additional averaging or increased photon counts. Instead, the SNR gain reflects PD’s ability to remove residual aberrations and redistribute signal into a sharper, higher-contrast PSF.

**Fig. 7.**
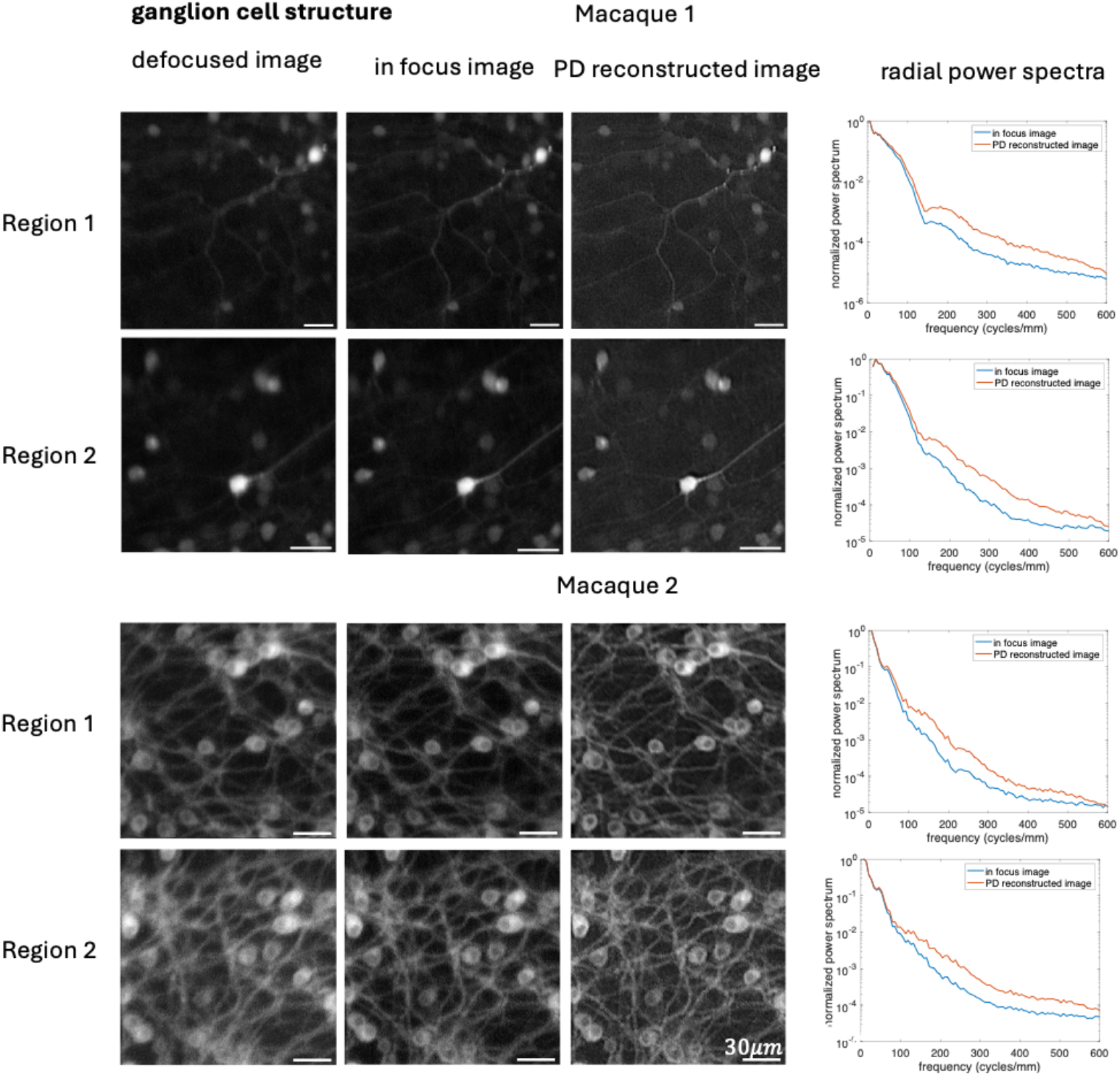
Fluorescence AOSLO images of retinal ganglion cells (RGCs) structures from two macaques, each with two retinal regions. For each region, the defocused image, in-focus image, and PD reconstructed image are shown side-by-side and in logarithmic scale. PD reconstruction enhanced the visualization of fine dendritic processes of RGC that were blurred or partially unresolved in the in-focus images. Radial power spectra (right) likewise show increased power at mid-to high-spatial frequencies following PD correction, consistent with improved recovery of structural detail. Scale bars: 30*μm*

**Fig. 8.**
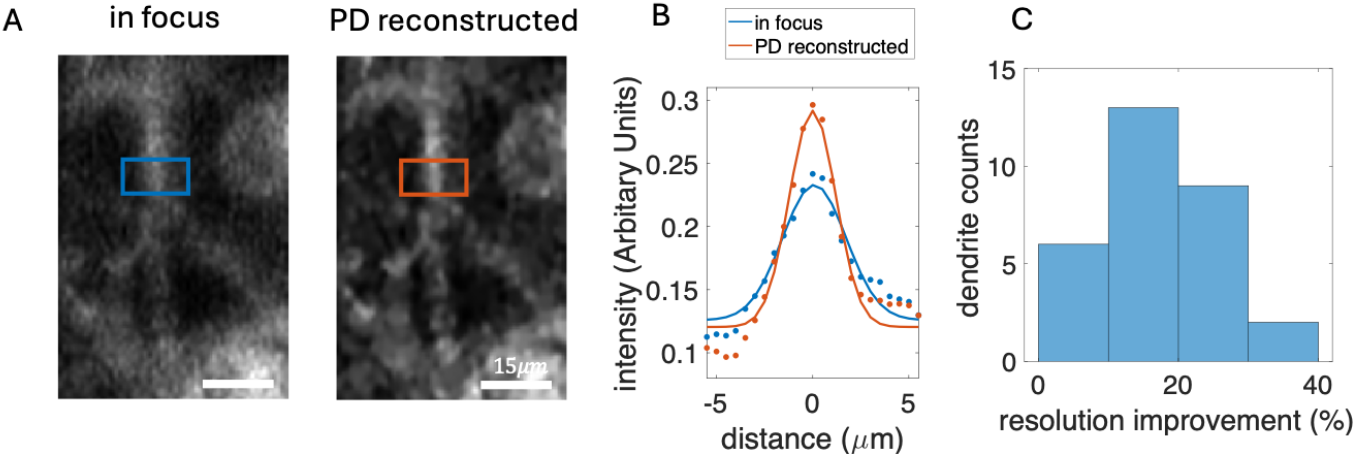
Improvement in resolution and signal-to-noise ratio (SNR) by PD reconstruction. (A) An example of comparison between the in-focus image and the PD-reconstructed image for fluorescently labeled dendrites. Scale bar: 5*μm*. (B) Intensity profiles across representative dendritic structures, highlighting increased peak intensity and reduced width in the PD-reconstructed image. (C) Histogram summarizing the resolution improvement across 30 analyzed dendritic structures, showing an average FWHM decrease of 18 ± 8%.

This improvement allowed clearer visualization of otherwise obscured dendritic structures, enabling more detailed biological interpretation of the underlying retinal morphology. The consistent enhancement across both cell types and imaging modalities demonstrated that PD can effectively compensate for residual aberrations left uncorrected by conventional AO, improving the resolution and signal-to-noise ratio (SNR) in dynamic *in vivo* primate retinal imaging.

## 4. Discussion

In this work, we demonstrated the feasibility of adapting phase diversity (PD) to confocal AOSLO for recovering residual aberrations beyond closed-loop AO correction. Our results showed that PD can improve the visualization of fine retinal structures, including cone mosaics and fluorescently labeled RGC dendrites, under *in vivo* imaging conditions. This expands the application of PD beyond its traditional use in astronomy and widefield microscopy, where it has been most thoroughly explored [10–16].

A key contribution of our study is tailoring the PD forward model to the confocal geometry of AOSLO and evaluating the effect of pinhole size on wavefront reconstruction. Unlike widefield systems, confocal AOSLO introduces unique challenges due to its localized illumination and spatial filtering through the pinhole. By explicitly modeling these effects, our implementation provides a framework for estimating aberrations in an imaging regime where standard PD formulations are not directly applicable. Our results revealed that the confocal pinhole size strongly influences the magnitude of aberrations that PD can recover. For a given amount of imposed aberration, smaller pinholes restrict detection to the central portion of the point spread function (PSF) for one scan position [30], causing light from aberrated regions outside the pinhole to be rejected. Consequently, PD tends to underestimate the total aberration magnitude when smaller pinholes are used, whereas larger pinholes capture the full extent of aberration-induced blur. Nevertheless, while smaller pinholes yield an underestimation relative to the introduced wavefront, they also provide intrinsically higher spatial resolution and are therefore more sensitive to residual aberrations that degrade image quality. In this sense, the underestimation obtained with smaller pinholes reflects the effective wavefront that governs image formation under confocal detection.

In addition to its advantages over hardware-based NCPA calibration, PD also differs fundamentally from conventional image deconvolution approaches, such as Wiener filtering [31], Richardson–Lucy (RL) [32,33] or blind deconvolution [34] which assume that the PSF is either known or can be estimated directly from the image. However, in confocal AOSLO the PSF depends on the wavefront aberration, the pinhole size, the excitation and detection geometry, and the confocal scanning process, making it poorly approximated by a fixed or shift-invariant kernel. PD uniquely addresses this by incorporating a parametric wavefront model and a deliberately introduced diversity phase, enabling simultaneous estimation of the object and the underlying aberration. The diversity image constrains the solution space and prevents confounding between object structure and optical blur—an ambiguity that classical deconvolution cannot resolve. As a result, PD retrieves the optically meaningful aberration that produced the confocal image, rather than simply sharpening the observed frame.

Despite the confocal pinhole size effects on PD reconstruction, PD is attractive because it requires only controlled phase diversity (e.g., defocus, astigmatism), works on the images acquired during normal operation, and can be run repeatedly to track changes. However, PD is not the only way to address residual aberrations, especially non-common path aberrations (NCPA). One direct calibration method bypasses the confocal aperture during a dedicated sequence, maximizes image sharpness, and derives the NCPA as the difference between the deformable mirror shapes that maximize sharpness and those minimizing the Shack– Hartmann metric [35]. This approach has the advantage of speed and robustness, and the correction can be incorporated into the AO loop as a static bias without sacrificing wavefront sensor bandwidth, but it requires separate calibration and does not capture aberrations that evolve during imaging. Simpler sensorless methods that maximize pinhole intensity with the confocal aperture in place have also been explored [36], but such strategies tend to favor throughput rather than resolution, meaning that brighter images do not necessarily correspond to sharper images in confocal AOSLO. In contrast, PD operates directly on acquired images, jointly recovering both the object and the aberration. This flexibility comes at the cost of requiring a second, deliberately degraded image, which in photon-limited applications, can reduces the effective signal-to-noise ratio because of the limited allowable exposure for light safety [37]. The benefits of improved aberration correction must therefore be weighed against the penalty in SNR.

While our work clearly demonstrated the utility of PD in enhancing confocal AOSLO images, several limitations merit discussion. First, the temporal separation inherent in acquiring sequential PD images introduces possible decorrelation caused by eye motion mainly due to heartbeat and respiration in the anesthetized primate, tear film changes or dynamic ocular aberrations within this time course [38,39]. Even with careful motion registration and averaging, such temporal variations can impose a limit on the achievable correction accuracy, particularly for higher-order modes.

Second, our implementation relied on a global estimation of the wavefront across the entire field of view (FOV). In our experiments, the FOV was restricted to less than 1.01°, under the assumption that residual aberrations remained approximately shift-invariant within this region. However, the eye’s optics, as well as small decentrations or tilts in the imaging beam, can introduce spatially varying aberrations outside the isoplanatic patch. The size of this patch can vary substantially across subjects, depending on individual ocular aberrations and alignment stability [40,41]. A natural extension of our work would be to separate images into smaller overlapping regions, applying PD locally to account for spatial variations in the wavefront. This could potentially recover fine-scale variations in residual aberrations, further improving local image quality for a larger FOV image.

Another possible development lies in integrating additional priors into the PD reconstruction. The retinal structures we aim to visualize, such as cone mosaics and dendritic processes, have characteristic spatial patterns [42,43]. Incorporating structural priors or regularization schemes that favor such biologically plausible features could help stabilize the inversion, especially in low signal-to-noise regimes [44].

## 5. Conclusion

In this study, we demonstrated that phase diversity (PD) can be effectively adapted to confocal adaptive optics scanning laser ophthalmoscopy (AOSLO) to correct residual aberrations that persist even after closed-loop AO correction. This approach enabled sharper visualization of cone mosaics and clearer delineation of fluorescently labeled RGC dendrites—features that are often degraded by residual aberrations. It opens new possibilities for non-invasive analysis of RGC morphology and the *in vivo* classification of distinct RGC types—information that previously required post-mortem histological examination. Nevertheless, applying PD in a confocal imaging configuration involves an intrinsic trade-off between optical resolution, optical sectioning, and the effective wavefront sensing, all of which are influenced by the confocal pinhole size.

Overall, our findings establish PD as a versatile computational supplement to conventional AO, capable of enhancing image contrast and resolution without requiring changes to the system. However, the practical implementation of PD does depend on the ability to introduce a known diversity phase. In our system, this is accomplished through the deformable mirror, which can apply the required phase on timescales of seconds. Other AOSLO platforms may or may not already have the necessary actuator bandwidth or software control to introduce the diversity phase during acquisition, and such considerations determine how readily PD can be integrated into different hardware configurations. Nonetheless, because the required phase change is small and deterministic, most existing ophthalmic AO systems with deformable mirrors or focus-tunable elements could, in principle, support PD acquisition with minimal modification. Although this work focused on AOSLO, the same framework can be extended to other adaptive optics modalities such as OCT and two-photon ophthalmoscopy.

## Acknowledgment

We thank Amber Walker and Jennifer LaPorta for assistance with animal anesthesia during imaging. We thank Qiang Yang for technical assistance. We also acknowledge the Penn Vector Core at the Perelman School of Medicine, University of Pennsylvania, and Shannon Boye and Sanford Boye at the University of Florida Vector Core Laboratory for viral vector production and support.

## Data availability

The data and code underlying the result section presented in this manuscript will be made publicly available upon publication at https://github.com/christiecai6/AOSLO-PhaseDiversity.git.

